# Contributions of Ccr4 and Gcn2 to the translational response of *C. neoformans* to host-relevant stressors and Integrated Stress Response induction

**DOI:** 10.1101/2022.07.26.501601

**Authors:** Corey M. Knowles, David Goich, Amanda L. M. Bloom, Murat C. Kalem, John C. Panepinto

**Author notes:** Address for Correspondence: John C. Panepinto, PhD, Professor of Microbiology and Immunology, Jacobs School of Medicine and Biomedical Sciences, 955 Main St, Ste 5229, Buffalo, NY 14203.

## Abstract

In response to the host environment, Cryptococcus neoformans must rapidly reprogram its translatome from one which promotes growth to one which is responsive to host temperature and oxidative stress. This reprogramming is primarily driven through the Gcn2-mediated repression of translation initiation and Ccr4-mediated removal of abundant pro-growth mRNAs from the translating pool. Here we investigate the contributions of these two pathways to the translational response to stress, and show that the response to oxidative stress is primarily driven by Gcn2 whereas temperature and oxidative stress both require Ccr4. Temperature stress, but not oxidative stress, result in an increase in RNase I resistant disome. Further, eIF2α phosphorylation varies in response to the type and magnitude of stress, yet all tested conditions induce translation of integrated stress response (ISR) transcription factor Gcn4, but not necessarily the Gcn4-dependent transcription. Finally, we define the ISR regulon in response to oxidative stress in C. neoformans. Together this study identifies the differential response to host-relevant stressors in an environmental fungus which can adapt to the environment inside the human host.

## INTRODUCTION

*Cryptococcus neoformans* is human-pathogenic fungus that, if not cleared from the lung, can cause cryptococcal meningoencephalitis, which kills nearly 180,000 people worldwide annually [1]. Upon inhalation by the human host, *C. neoformans* is suddenly subjected to a multitude of stressors, including but not limited to the mammalian core temperature of 37°C and oxidative stress produced by macrophages in the lungs. To adapt to and survive the stress from such an extreme environmental transition, *C. neoformans* reprograms its translatome, translating mRNAs coding for effectors of stress mitigation rather than those that promote cell growth [2–5].

Translatome reprogramming is comprised of two molecular processes. First, the removal of abundant mRNAs that are abundant and efficiently translated, such as ribosomal protein (RP) mRNAs, freeing resources for the translation of stress-responsive mRNAs [3,6]. Second, the initiation of translation on new mRNAs entering the translating pool, which is governed by the availability of the ternary ribonucleoprotein complex, in which the initiator methionyl-tRNA and eIF2-GTP bind to the 40S ribosome to form the 43S pre-initiation complex (reviewed in [7]).

The removal of mRNAs from the translating pool occurs primarily through coordinated mRNA decay, for which the initial and rate-limiting step is deadenylation, performed by Ccr4 in *C. neoformans* [8]. Under stress, the alpha subunit of eIF2 is phosphorylated by Gcn2, the sole eIF2α kinase in this organism, which represses translation initiation[5]. When both translation initiation and programmed mRNA decay occur simultaneously, the translatome is reprogrammed for the production of stress-responsive effectors [6].

An additional consequence of eIF2α phosphorylation is the induction of the integrated stress response (ISR) [9,10]. When eIF2α is phosphorylated, the 43S preinitiation complex bypasses upstream open reading frames of the mRNA encoding the transcription factor Gcn4 that typically repress its translation [11]. Newly translated Gcn4 then translocates to the nucleus, activating the transcription of stress-responsive mRNAs [12]. Gcn4 targets, which comprise the ISR regulon in fungi, are then introduced into the translating pool, facilitating stress adaptation. The ISR regulon of *C. neoformans* remains undefined at this time.

A growing body of evidence has begun to uncover how environmental stress is sensed by cells. Osmotic stress, heat shock, and stress from translation inhibitors, RNA-damaging chemicals, tRNA availability, and mRNA defects have all been shown to result in stalled or collided ribosomes [13–20]. Stalled or collided ribosomes serve as a trigger and signaling platform for various downstream events, which include Gcn2 activation [21–25] and subsequent ISR induction, ribosome quality control, nascent polypeptide degradation, and mRNA surveillance and decay [19,26–36]. However, ribosome collisions and the downstream consequences in fungal pathogens have yet to be described. In this study, we sought to determine which stressors lead to translational stress-induced ribosome collision and characterize the contribution from both regulation of mRNA entry and exit of the translating pool on translatome reprogramming in *C. neoformans*. We assess host-relevant stressors, and determine which stressors lead to translational stress-induced ribosome collision. In addition, we examine how stress-induced translatome regulation induces the ISR and define the ISR regulon in *C. neoformans*.

## METHODS AND MATERIALS

### Strains

*C. neoformans* var. *grubii* H99 serotype A (taxid:235443) was used as the WT strain in this study. All mutant strains were created in this WT background. The *ccr4*Δ strain used was previously published [2]. Other mutant strains were constructed using methods previously described [37]. To make the *ccr4*Δ::*gcn2*Δ strain, a *gcn2*Δ strain was first constructed as described previously [5], with the exception that the homologous arms were flanked by a G418 resistance cassette amplified with primers F-NEO-*Bg*/II and R-NEO-*Sac*I listed in Table S1 and used to transform the WT strain. The *gcn2*Δ strain was then transformed with the *CCR4* knockout construct previously published [2]. The *CCR4*::*ccr4*Δ strain was constructed by amplifying the *CCR4* genomic locus from the WT strain using primers F-CCR4-1kbUp-Spe1 and R-CCR4- 1kbDown-Spe1 (Table S1) and inserting it into the *ccr4*Δ mutant [2]. All strains were confirmed by PCR, Northern blotting, and phenotypic profiling.

### Media and growth conditions

All experimental cultures were started from overnight cultures grown in YPD (1% yeast extract, 2% peptone, and 2% dextrose) at 30°C. Experimental cultures were seeded to an optical density at 600 nm of 0.18–0.20 in YPD (unless otherwise noted). YNB (BD Difco Cat#: 291920) supplemented with 2% dextrose was used as the minimal medium. Cultures were grown at 30°C until the mid-logarithmic phase was reached. Cells were pelleted at 3,000 rcf for 2 min and resuspend in prewarmed 37°C medium for temperature stress or with medium containing 1 mM or 2 mM H_2_O_2_ (Fisher Chemical, Cat#:H325-500) for oxidative stress or 40 mM 3-AT (TCI Cat#: A0432). Cultures were incubated for the times indicated in the text and figures prior to harvesting. Cells were harvested by pelleting for 2 min at 3,000 rcf and were then flash frozen in liquid nitrogen.

### Spot plating

Strains were grown in YPD overnight at 30°C. The following day, cells were pelleted at 3,000 rcf for 2 min and washed twice in sterile deionized water. The optical density at 600 nm of each culture was set to 1.0 in sterile deionized water, and the cultures were then serially diluted 10-fold six times. Five microliters was then spotted onto either YPD or YNB+2% dextrose agar, with or without H_2_O_2_, and incubated at 30°C, 37°C, or 38°C for 72 h.

### Polysome and disome profiling

Each cell pellet was resuspended in 1 mL of polysome lysis buffer (20 mM Tris-HCl [pH 8.0], 140 mM KCl, 5 mM MgCl_2_, 1% Triton X-100, 25 mg/mL heparin sodium sulfate, and 0.1 mg/mL cycloheximide) and transferred to a microcentrifuge tube. Cells were repelleted by centrifugation at 2,350 rcf for 2 min, resuspended in 50 μL polysome lysis buffer, and layered on glass beads for lysis using a Bullet Blender cooled with dry ice (5 min at setting 12). An additional 150 μL of lysis buffer was added to the glass beads. Lysate was removed from the glass beads and centrifuged at 20,050 rcf for 10 min. The cleared cell lysate was removed from the pellet, and RNA was quantified using a NanoDrop (Thermo Fisher). For each profile, 250 μg of RNA was loaded onto a 10–40% sucrose gradient and centrifuged at 260,800 rcf for 120 min in a SW41Ti swinging bucket rotor. A Brandel tube piercer was used for the determination of RNA absorbance of the gradients at 254 nm with a Teledyne UA-6 detector. Data were captured using a DATAQ DI-1110 and recorded using DATAQ WinDaq software.

### Western blotting

Pelleted cells were thawed and resuspended in 1 mL of sterile deionized water and transferred to 1.5 mL microcentrifuge tubes. Cells were again pelleted by centrifugation at 2,400 rcf for 2 min and resuspended in 50 μL of lysis buffer (10 mM HEPES, 100 mM KCl, 5 mM MgCl_2_, 0.5% NP-40, 10 μM dithiothreitol, and 10 μL/mL Halt protease inhibitor). Cells were layered onto glass beads and lysed using a Bullet Blender as described above. Lysate was washed from the glass beads using an additional 30 μL of lysis buffer. The whole-cell lysate was then cleared by centrifugation at 21,000 rcf for 10 min. Protein was quantified using a Pierce 660-nm protein assay kit, and equivalent amounts of protein from each sample were boiled in Laemmli buffer containing 2- mercaptoethanol before loading onto Bio-Rad stain-free Tris-glycine gels for electrophoretic separation. Total protein was quantified by imaging the stain-free gels using a Bio-Rad Gel Doc XR+ and Image Lab software. Protein was transferred to polyvinylidene difluoride membranes using a Bio-Rad Trans-blot Turbo, and membranes were blocked with Bio-Rad EveryBlot blocking buffer. Blots were probed for phosphorylated eIF2α using anti-eIF2α (phospho-S51) antibody (Abcam, Cat#: ab32157, 1: 1000 dilution). Blots were probed for Gcn4 using a polyclonal rabbit anti-Gcn4 antibody raised against recombinant *C. neoformans* Gcn4 by GenScript. The secondary antibody used was horseradish peroxidase-conjugated anti-rabbit IgG (Cell Signaling Technologies, Cat#: 7074, 1:10,000 dilution). Horseradish peroxidase was visualized via Bio-Rad Clarity Max (Cat#: 1705062) and imaged on a Bio-Rad ChemiDoc MR. Quantification of blots was performed in Bio-Rad Image Lab.

### Northern blotting

Cells were thawed and resuspended in Qiagen RLT buffer containing 10 μL/mL 2- mercaptoethanol. The lysates were layered in glass beads and mechanically lysed using a Bullet Blender as described above. Lysates were flushed from the glass beads using the same buffer, and whole-cell lysates were cleared by centrifugation at 21,000 rcf for 10 min. The supernatants were removed, and equal volumes of 70% ethanol were added to precipitate nucleic acids. RNA was then isolated using the Qiagen RNeasy kit according to the manufacturer’s instructions. RNA was quantified using a NanoDrop, and 5 μg was loaded onto an agarose-formaldehyde gel for electrophoretic separation. Total RNA was imaged using a Bio-Rad Gel Doc and ImageLab software. RNA was transferred to a nylon membrane and probed using a ^32^P-labeled DNA probe, created using Invitrogen RadPrime DNA Labeling kit (Cat#: 18428011) as described previously [38]. The *RPL2* probe was made as described previously [38]. The *ARG1* probe was created using genomic DNA amplified using ARG1-Northern-F and ARG1-Northern-R primers listed in Table S1. Blots were exposed to a phosphor screen, and the screen was scanned using a GE Typhoon phosphoimager. Signals were quantified using Quantity One software.

### RNA sequencing and analysis

WT and *gcn4*Δ strains were grown to mid-logarithmic phase at 30°C, and cells were treated with 1 mM H_2_O_2_, or left untreated, for 1 h. RNA was extracted as described for Northern blotting and treated using an on-column DNase kit (Qiagen). RNA quality was determined by investigating integrity on an RNA gel prior to sequencing. RNA samples were submitted to Genewiz (Azenta) for poly(A)+ purification, library preparation, and Illumina sequencing. The sequencing data can be viewed at GEO under accession number: **GSE206508.** Two biological replicates were analyzed per strain. RNA sequencing reads were trimmed to remove adapter sequences using Cutadapt [39]. Filtered reads were aligned to the *C. neoformans* H99 genome (FungiDB) using STAR alignment [40]. Read counts for each gene were calculated using RSEM [41], and differentially expressed genes across samples were determined using DESeq2 in R [42]. Data were then filtered according to a 1.75-fold change and an adjusted *P* value threshold of ≤ 0.05.

The ISR was defined by the following three criteria: (i) genes that were downregulated in the *gcn4*Δ strain treated with 1 mM H_2_O_2_ compared to expression in the WT treated with 1 mM H_2_O_2_, (ii) genes that were upregulated in the WT upon treatment with 1 mM H_2_O_2_, and (iii) genes that were not upregulated in the *gcn4*Δ mutant upon treatment with 1 mM H_2_O_2_. Differential expression of genes was visualized using the R-package EnhancedVolcano [43], with ISR genes highlighted in green and the ribosome biogenesis GO term (GO:0042254) highlighted in red. GO analysis of the ISR genes was performed using FungiDB, and visualized using the R package ggplot2 [44].

## RESULTS

### *C. neoformans* exhibits ribosome collision and translational repression in response to temperature and oxidative stresses

We first set out to compare the translational responses of wild-type (WT) *C. neoformans* to temperature and oxidative stress by rapidly shifting the temperature of liquid cultures from 30°C to 37°C and by treating them with H_2_O_2_, respectively. To isolate the effects of environmental stressors, we included samples treated with 40 mM 3-amino-1,2,4- triazole (3-AT), a well-characterized inhibitor of translation. 3-AT causes ribosomes to pause at histidine codons as a result of reduced histidine biosynthesis thereby reducing hystidyl-tRNA levels in the cell, and results in ribosome pausing at histidine codons [45]. The combination of these events results in Gcn2-mediated phosphorylation of eIF2α.

Western blotting revealed a modest and transient phosphorylation of eIF2α in response to the switch to 37°C (**Fig 1A**). The phosphorylation was more pronounced in response to 1 mM H_2_O_2_, but 2 mM H_2_O_2_ induced a strong response that was sustained for at least 60 min. Sustained phosphorylation was also achieved with 40 mM 3-AT, though the levels of phospho-eIF2α were more similar to those observed with 1 mM H_2_O_2_.

**Fig 1.**
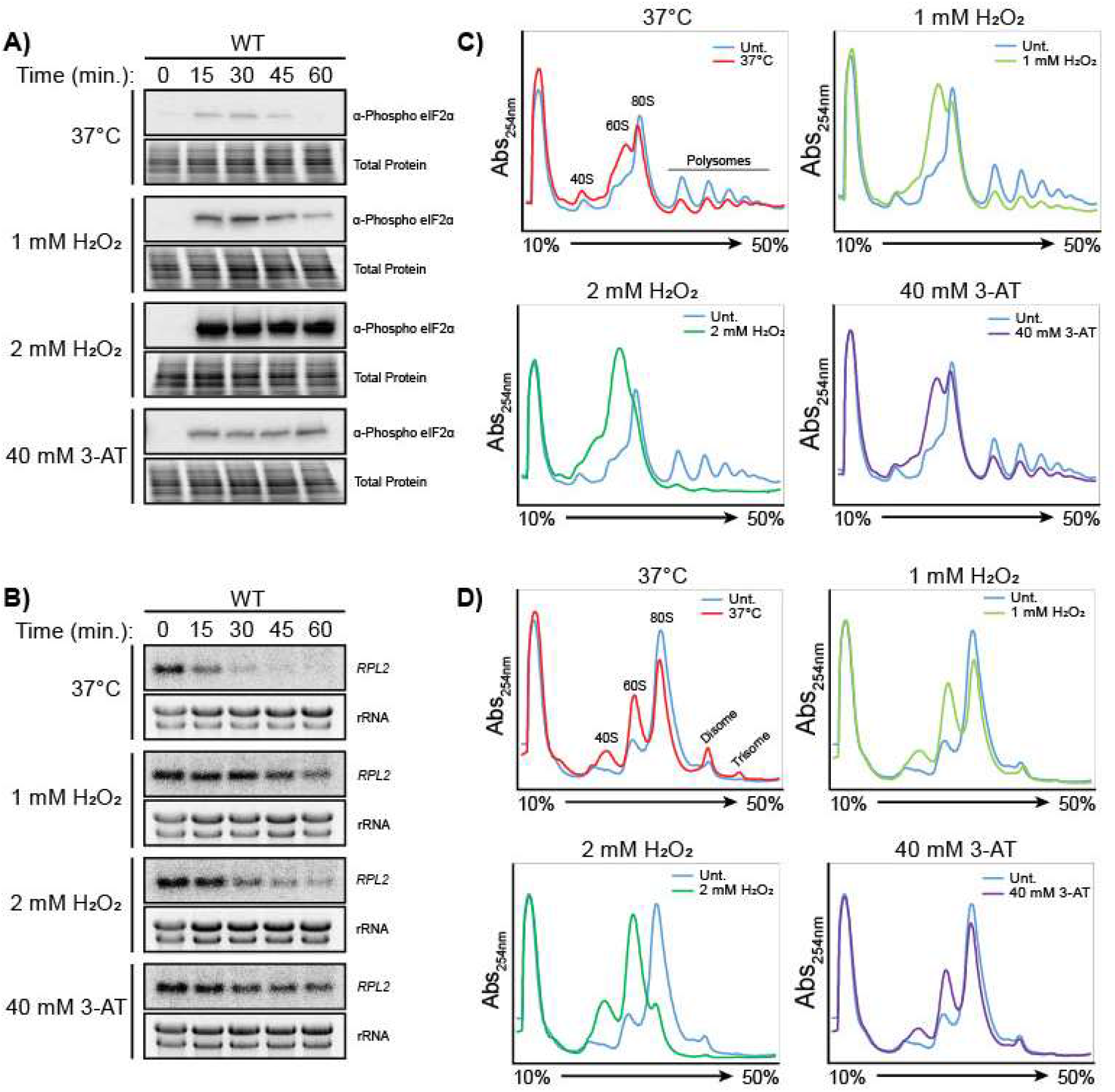
*C. neoformans* represses translation in response to temperature and oxidative stresses. Western blots for phosphorylated eIF2α (**A**) and Northern blots for the *RPL2* transcript (**B**) in *C. neoformans* in response to a shift to 37°C, 1 mM H_2_O_2_, 2 mM H_2_O_2_, and 40 mM 3-AT. (**C**) Polysome profiling of *C. neoformans* in response to a shift to 37°C, 1 mM H_2_O_2_, 2 mM H_2_O_2_, and 40 mM 3-AT. (**D**) Polysome profiling analysis of samples from panel C after treatment with RNase I.

We previously showed that *C. neoformans* reduces levels of ribosomal protein (RP) transcripts in response to various stressors and that the reduction in response to 37°C stress is necessary for stress-responsive translatome reprogramming [5,46–48]. Therefore, we measured the levels of RP transcript *RPL2* in response to temperature, oxidative stress, and 3-AT treatment. As expected, 37°C stress rapidly reduced *RPL2* levels (**Fig 1B**). Oxidative stress also reduced *RPL2* levels, with a greater effect from 2 mM H_2_O_2_ than from 1 mM H_2_O_2_. A mild reduction in *RPL2* was observed in the 3-AT-treated samples, suggesting that mechanisms other than Gcn2 activation contribute to the reductions seen after temperature or oxidative stress.

We performed polysome profiling to compare the overall translational states in *C. neoformans* exposed to temperature and oxidative stresses. The stress conditions of 37°C and 1 mM H_2_O_2_ resulted in a moderate collapse of the polysomes, similar in magnitude to the collapse seen with 3-AT treatment (**Fig 1C**). Specifically, there was an increase in the 60S peak, with only a minor decrease in the 80S peak, indicative of repressed translation initiation. Under stress from 2 mM H_2_O_2_, there was a greater magnitude of polysome collapse, with a greater increase in the 60S peak, which eclipsed the 80S peak. This suggests a much greater level of translational repression than that observed in response to the other three tested conditions.

To determine if the stressors would also induce ribosome collision, we performed disome profiling, in which disomes are visualized as a peak resistant to collapse following RNase I digestion. Disome profiles were captured after 30 min of stress treatment, corresponding to the time when we saw the greatest levels of eIF2α phosphorylation from temperature and 1 mM H_2_O_2_ stress. Surprisingly, an increase in disomes and trisomes was observed in response to 37°C stress, the stress condition which exhibited the smallest increase in eIF2α phosphorylation levels (**Fig 1D**). There was no change in disome accumulation as a result of 1 mM H_2_O_2_ or 3-AT treatment, and there was a decrease in the quantity of disomes in the 2 mM H_2_O_2_-treated samples.

Together, these data suggest that although both temperature and oxidative stress result in translational repression (as observed by eIF2α phosphorylation, reduced *RPL2* levels, and polysome collapse), the mechanisms and degrees of translational repression vary with the type and magnitude of the stress. These data suggest that a feedback mechanism may be functioning where increased eIF2α phosphorylation may serve to limit the number of ribosomes on mRNAs as the cells respond to stress. Additionally, 1 mM H_2_O_2_ stress may result in individually stalled ribosomes which are unable to collide and are thus unable to be detected using an RNase I protection assay.

### Gcn2 is required for translational repression during oxidative stress but is dispensable during temperature stress

To determine the contribution of Gcn2 to the stress adaptations we observed in *C. neoformans,* we compared the responses to both temperature and oxidative stresses to those in a *gcn2*Δ mutant strain that is unable to phosphorylate eIF2α. We performed serial dilution spot assays, subjecting the strains to the temperature stress, oxidative stress, and combinations of temperature and oxidative stress, which more closely mimic the conditions that occur during the course of an infection. Although we expected the *gcn2*Δ mutant to exhibit temperature sensitivity given the observed eIF2α phosphorylation and increase in 60S peak in the WT strain during temperature stress (**Fig 1A, C**), the growth of the *gcn2*Δ mutant did not differ from that of the WT at 37°C or 38°C (**Fig 2A**). The *gcn2*Δ strain showed sensitivity to 3 mM H_2_O_2_ at 30°C. Interestingly, the growth of the *gcn2*Δ strain was severely stunted when oxidative stress and temperature stress were combined. These growth phenotypes were restored when the gcn2Δ mutant was complemented with the WT *GCN2* gene (**Fig 2A**). These data suggest that the two stressors are not only sensed differently but also additive and that Gcn2 activity becomes more important under cumulative stress.

**Fig 2.**
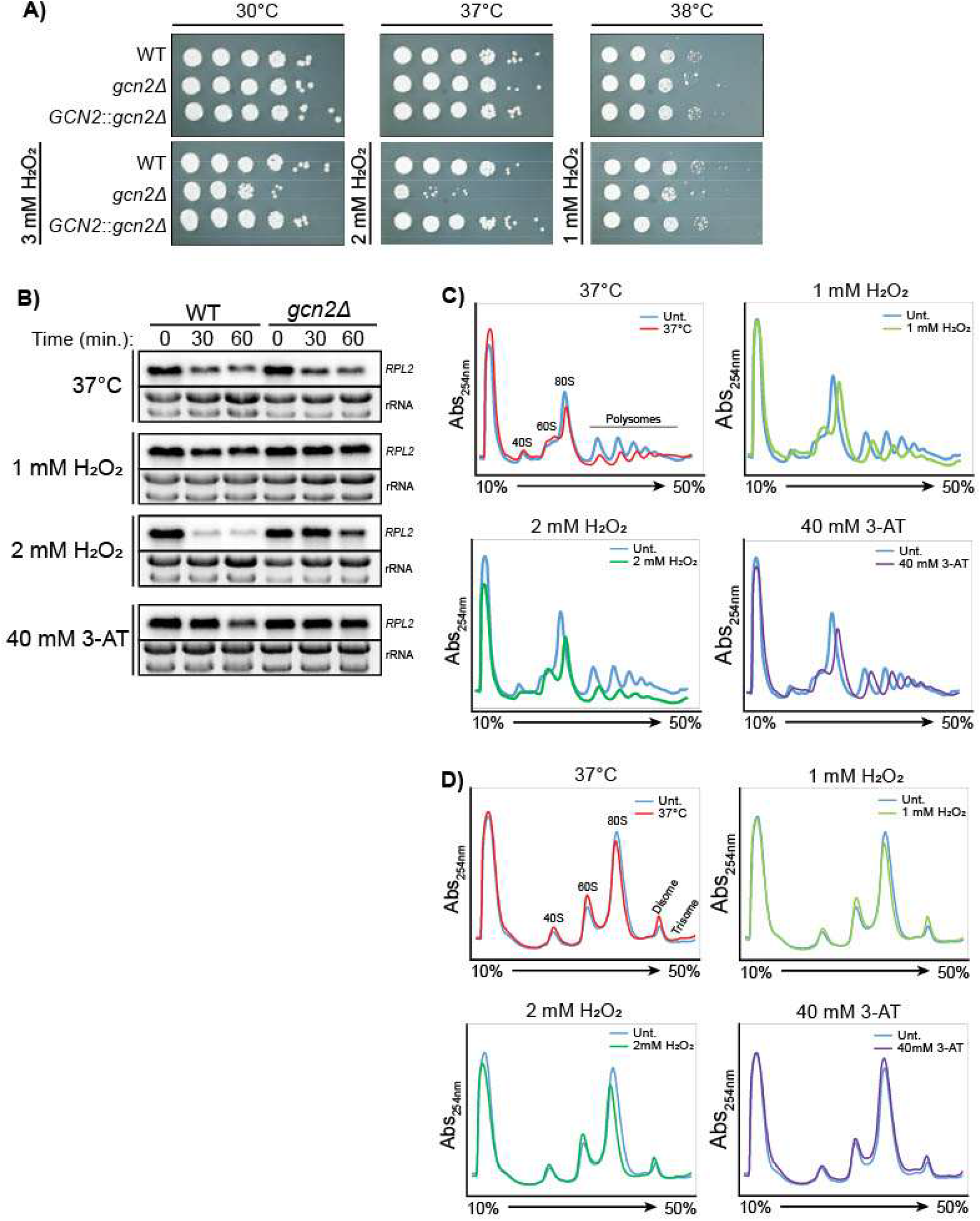
Gcn2 is primarily required for translational repression under oxidative stress. (**A**) Serial dilution spot plate analysis of WT, *gcn2*Δ mutant, and *GCN2*::*gcn2*Δ complemented strains. (**B**) Northern blot analysis for the *RPL2* transcript in WT and *gcn2*Δ mutant strains in response to 37°C, 1 mM H_2_O_2_, 2 mM H_2_O_2_, and 40 mM 3-AT. (**C**) Polysome profiling of the *gcn2*Δ mutant in response to 30 min at 37°C or 1 mM H_2_O_2_, 2 mM H_2_O_2_, and 40 mM 3-AT. (**D**) Polysome profiling analysis of RNase I-digested samples from panel C.

Northern blots showed that *RPL2* levels were suppressed similarly in the WT and *gcn2*Δ strains under 37°C stress (**Fig 2B**). However, the *gcn2*Δ mutant failed to reduce *RPL2* levels under oxidative stress with 1 and 2 mM H_2_O_2_, unlike the WT strain. Similar results were obtained with 3-AT treatment; whereas the WT showed a moderate reduction in *RPL2* at 60 min, no reduction was detectable in the *gcn2*Δ mutant.

Polysome profiling confirmed that the *gcn2*Δ mutant is defective in polysome repression in response oxidative stress, congruent with previously reported data [5]. In contrast to that for the WT strain (**Fig 1C**), no increases in 60S peaks and reduced collapse of polysome peaks were observed for the mutant in response to 1 mM or 2 mM H_2_O_2_ (**Fig 2C**). 3-AT treatment similarly had no effect on the ribosome profile of the *gcn2*Δ mutant.

Because Gcn2 activity was recently linked to ribosome collision and subsequent quality control events [21], we hypothesized that ribosome collisions would accumulate in a strain which may be defective in recognizing stalled/collided ribosomes or transducing subsequent signaling events. Disome profiling in the *gcn2*Δ mutant after a 30-min treatment demonstrated only a modest increase in disomes, which was observed under all stress and treatment conditions (**Fig 2D**). Importantly, there was an retention of disomes when the *gcn2*Δ strain was treated with 2mM H_2_O_2_, unlike in the WT strain, which may be a result of a failure to effectively repress entry of oxidatively damaged mRNAs into the translating pool.

These data indicate that Gcn2 is required for *RPL2* repression and polysome collapse during oxidative stress but is dispensable during temperature stress. However, Gcn2 improves survival under temperature stress when other stressors are present. Additionally, Gcn2 dependent regulation of translation initiation contributes to the prevention and/or resolution of ribosome collisions that occur in response to severe oxidative stress.

### Ccr4 is required for the translational response to temperature and oxidative stresses

A *ccr4*Δ *C. neoformans* mutant deficient in deadenlylation-dependent mRNA decay has severely attenuated virulence in a mouse model and is broadly stress sensitive, suggesting that Ccr4 contributes to stress adaptation in general [49]. We previously showed that the *ccr4*Δ mutant is defective in translatome reprogramming in response to temperature stress [2,4]. Therefore, we assessed if this defect occurs in response to other physiologically relevant stressors, namely, oxidative stress alone and in combination with temperature stress. The results from a spot plate assay demonstrated that, in addition to temperature sensitivity, the growth of the *ccr4*Δ mutant was also sensitive to oxidative stress at 30°C (**Fig 3A**). Moreover, the defect was more pronounced under compound stress conditions of 37°C with 2 mM H_2_O_2_. and 38°C with 1 mM H_2_O_2_. All growth phenotypes were restored when the *ccr4*Δ mutant was complemented with the WT *CCR4* gene (**Fig 3A**).

**Fig 3.**
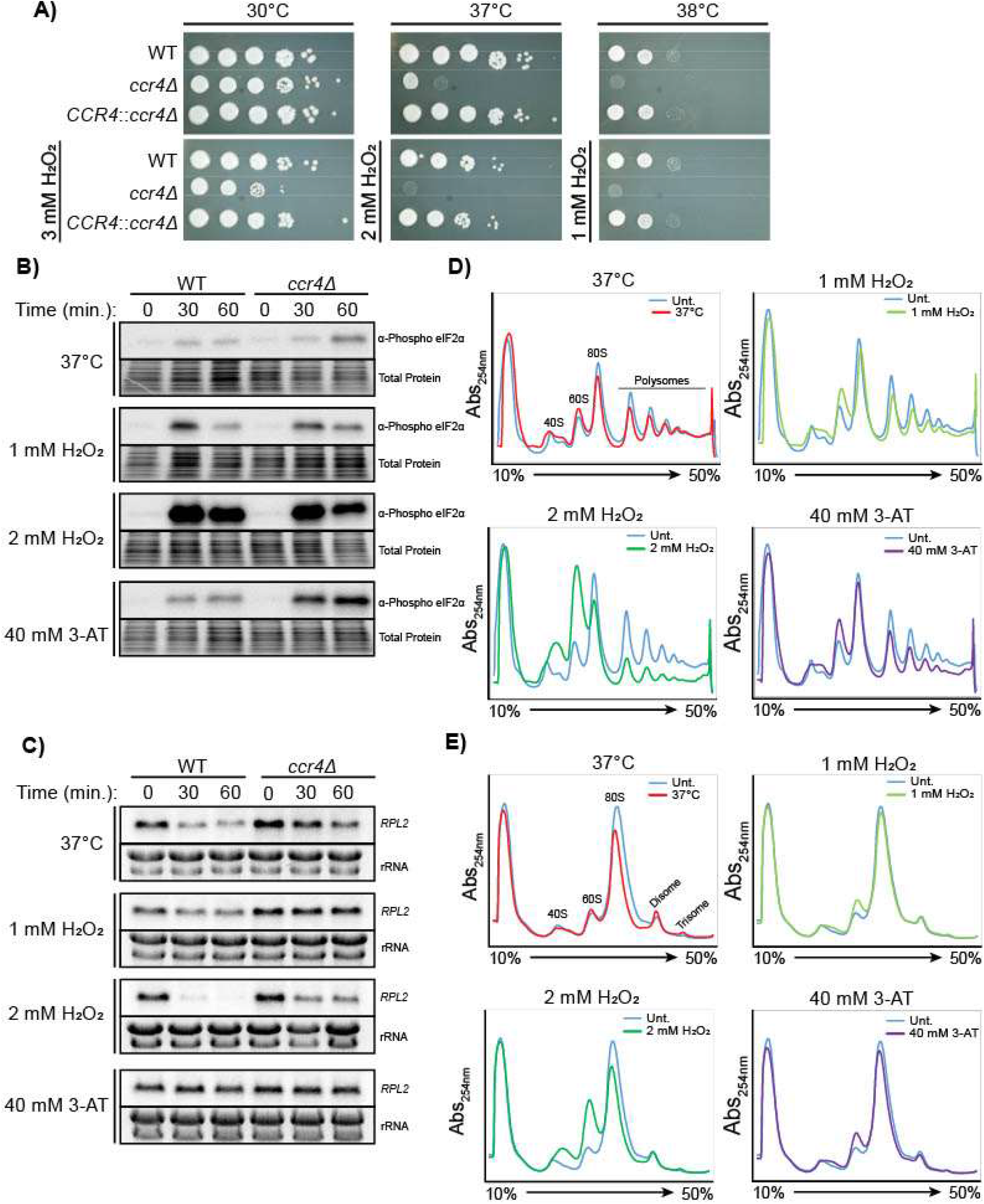
Ccr4 is required for translational repression in response to temperature and oxidative stresses. (**A**) Serial dilution spot plate analysis of WT, *ccr4*Δ mutant, and *CCR4*::*ccr4*Δ complemented strains. Western blots for phosphorylated eIF2α (**B**) and Northern blots for the *RPL2* transcript (**C**) in WT and *ccr4*Δ strains in response to 37°C, 1 mM H_2_O_2_, 2 mM H_2_O_2_, and 40 mM 3-AT. (**D**) Polysome profiling of the *ccr4*Δ mutant after 30 min at 37°C or 1 mM H_2_O_2_, 2 mM H_2_O_2_, and 40 mM 3-AT. (**E**) Polysome profiling analysis of RNase I-digested samples from panel D.

We then asked if translation initiation was repressed in *ccr4*Δ to compensate for the retention of mRNAs in the translating pool [3]. Western blot analysis showed that eIF2α phosphorylation was delayed but increased in a *ccr4*Δ mutant relative to the WT under 37°C stress, and was overall increased in response to 3-AT treatment (**Fig 3B**). Under both oxidative stress conditions eIF2α phosphorylation was unchanged in the *ccr4*Δ mutant relative to the WT (**Fig 3B**). Loss of Ccr4 also led to relative protection of *RPL2* transcripts in response to temperature and oxidative stresses as well as 3-AT treatment (**Fig 3C**), suggesting that Ccr4 universally promotes RP mRNA decay.

Polysome profiling showed that, unlike the WT strain (**Fig 1C**), the *ccr4*Δ strain retained low-molecular-weight polysomes and failed to accumulate high-molecular-weight polysomes under 37°C stress (**Fig 3D**), consistent with its previously reported translatome reprogramming defect [3]. No increase in the 60S peak was observed, also consistent with previous reports [3]. Furthermore, the *ccr4*Δ strain failed to show polysome collapse or increases in monosome and 60S peaks in response to 1 mM H_2_O_2_ that were noted in the WT strain (compare **Fig 3D** and **Fig 1C**). In response to 2 mM H_2_O_2_, the *ccr4*Δ mutant exhibited polysome and 80S peak collapse with an increase in subunit peaks, but not to the extent seen in the WT strain. The polysome collapse in the *ccr4*Δ mutant in response to 3-AT treatment was similar to that observed in the WT strain; however, the 60S peak did not increase to the same extent.

Surprisingly, disomes accumulated in the *ccr4*Δ mutant to a similar extent as in the WT strain under untreated, temperature, and 1 mM H_2_O_2_ conditions (**Fig 3E**), suggesting that Ccr4 is not involved in clearing collided ribosomes. Unlike that in the WT strain, the disome peak was retained in the *ccr4*Δ strain after treatment with 2 mM H_2_O_2_. This suggests that the cell’s response to 2 mM H_2_O_2_ is distinct from that for the other three stressors tested, and that a defect in translational repression may result in perpetuation of ribosome collision.

We conclude from these data that the *ccr4*Δ mutant is defective in eliminating abundant RP transcripts (e.g., *RPL2*) under temperature and oxidative stresses, resulting in an increased sensitivity to these stressors and a reduction in fitness. Thus, deadenylation-dependent mRNA decay impacts translatome reprogramming in response to both temperature and oxidative stress, whereas regulation of translation initiation thus far appears only to impact the translational response to oxidative stress.

### Additional stress from growth in minimal medium contributes to the translational response

So far, the experiments assessing translatome reprogramming were performed in nutrient-rich yeast extract-peptone-dextrose (YPD) medium. We reasoned that the additional stress of growth in a minimal medium (with only dextrose and ammonium as sources of carbon and nitrogen, respectively), which more closely mimics the nutritional environments seen inside the human host, would further impact translational capacity. Thus, we compared *C. neoformans* strains grown in YPD to those grown in yeast nitrogen base (YNB) with dextrose. The *ccr4*Δ strain grew better on YNB than on YPD at 37°C (**Fig 4A**). This was also observed in the WT strain at 38°C, although to a lesser extent (**Fig 4A**). However, the protection afforded by minimal medium (i.e., YNB) was lost under conditions of oxidative stress, i.e., when 1 mM H_2_O_2_ was added.

**Fig 4.**
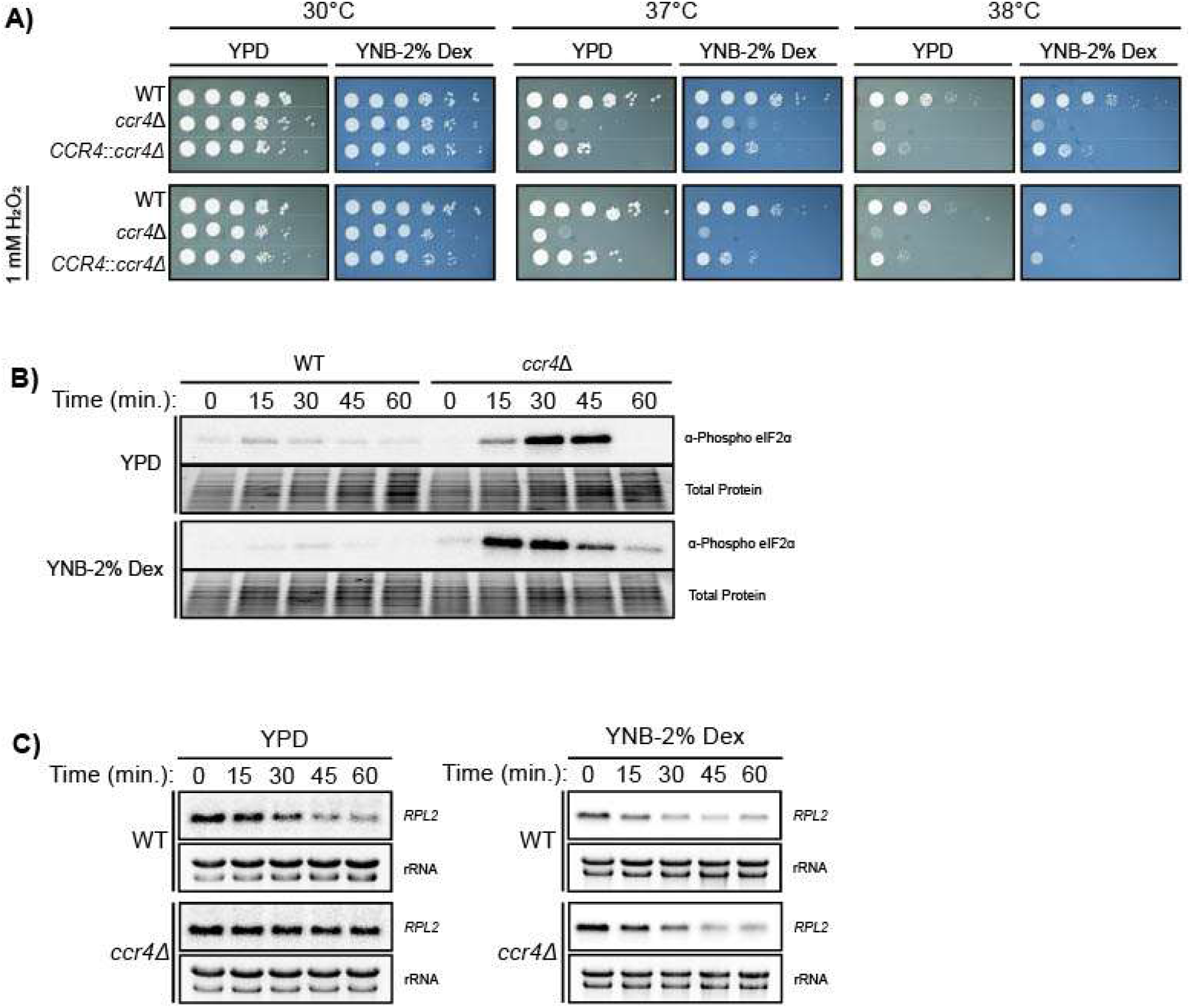
Minimal medium is a stressful environment that results in increased repression of translation initiation. (**A**) Serial dilution spot plate analysis of WT, *ccr4*Δ mutant, and *CCR4*::*ccr4*Δ complemented strains. (**B**) Western blots for phosphorylated eIF2α in WT and *ccr4*Δ mutant strains in response to 37°C in nutrient-rich medium (YPD) and minimal medium (YNB-2% dextrose). (**C**) Northern blot analysis for the *RPL2* transcript in WT and *ccr4*Δ mutant strains in response to 37°C in either YPD or YNB-2% dextrose.

The phosphorylation of eIF2α in response to 37°C stress was initiated sooner in the *ccr4*Δ strain grown in minimal medium (**Fig 4B**). We then investigated if this enhancement affected the clearance of *RPL2* from the translating pool. Indeed, the levels of *RPL2* were reduced in the *ccr4*Δ strain when it was grown in YNB medium, resulting in a reduction in *RPL2* transcripts that paralleled that in the WT (**Fig 4C**).

These data suggest that increased eIF2α phosphorylation levels, or otherwise altered translation kinetics in minimal media, contribute to translatome reprogramming in the absence of Ccr4-mediated clearance of RP transcripts, conferring some degree of protection.

### Gcn2 can compensate for the loss of Ccr4 in the translational response to temperature in minimal medium

The role of Gcn2 in the effects of YNB on the growth and translational responses of the *ccr4*Δ strain were examined by generating a *ccr4*Δ::*gcn2*Δ double mutant. Growth on YNB medium afforded the *ccr4*Δ::*gcn2*Δ mutant some protection against 37°C stress, similar to that observed for the *ccr4*Δ mutant, but not against oxidative stress (**Fig 5A**). The combinatorial stress of 37°C and 1 mM H_2_O_2_ synergistically affected growth of the *ccr4*Δ::*gcn2*Δ mutant, regardless of the medium, which is in agreement with the data showing that both Ccr4 (**Fig 3A**) and Gcn2 (**Fig 2A**) contribute to fitness under oxidative stress. However, the *ccr4*Δ::*gcn2*Δ double mutant did not differ from the *ccr4*Δ mutant when under the combinatorial stress, also consistent with the data showing that Gcn2 does not contribute to the temperature stress response (**Fig 2A**).

**Fig 5.**
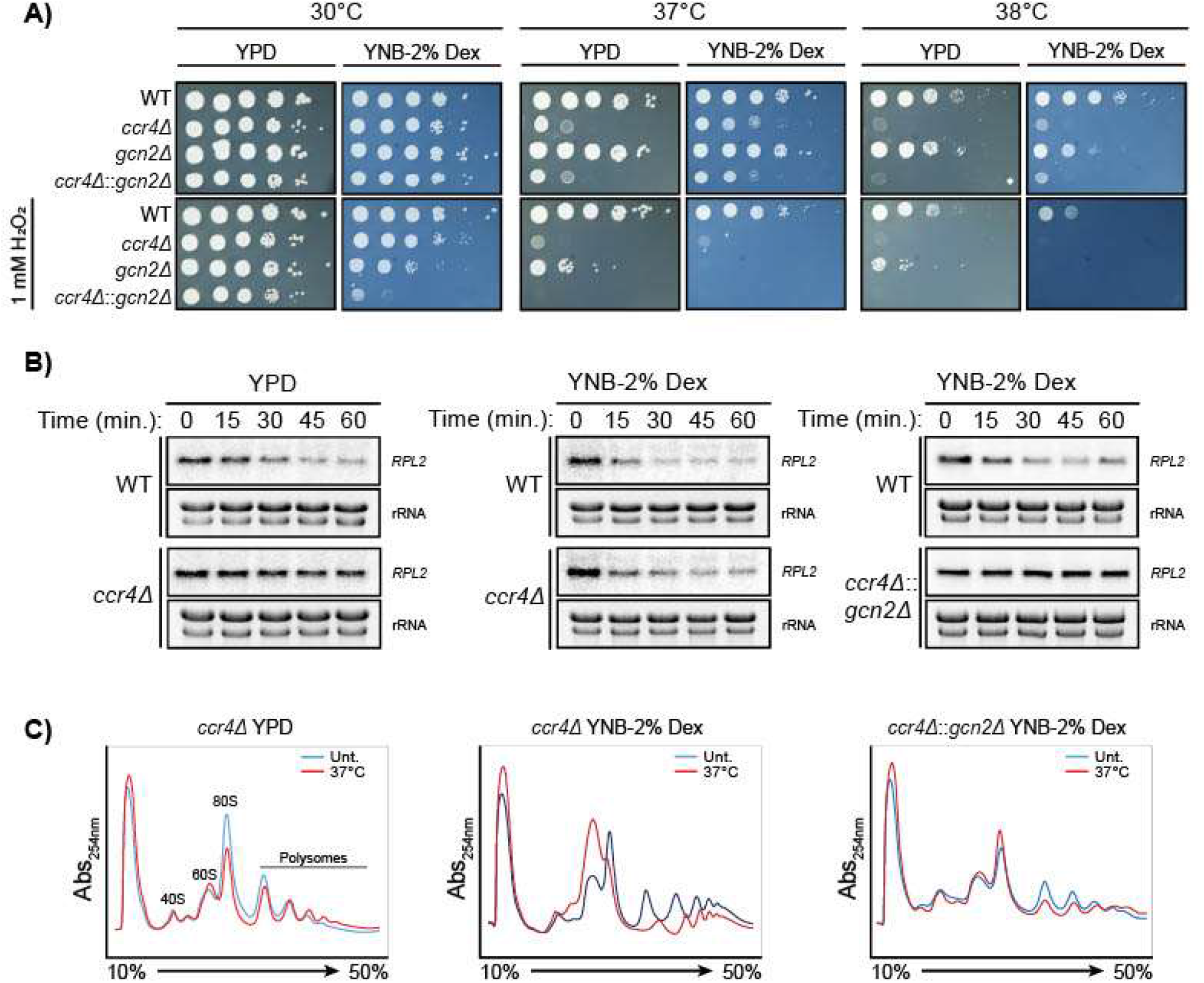
Gcn2-mediated translational repression in minimal medium can compensate for an absence of mRNA during translatome reprogramming. (**A**) Serial dilution spot plate analysis of WT, *ccr4*Δ, *gcn2*Δ, and *ccr4*Δ::*gcn2*Δ mutant strains at different temperatures and in the presence of 1 mM H_2_O_2_ in YPD or YNB-2% dextrose. (**B**) Northern blot analysis for the *RPL2* transcript in WT and *ccr4*Δ or ccr4Δ::gcn2Δ strains in response to 37°C in either YPD or YNB-2% dextrose. (**C**) Polysome profiling of *ccr4*Δ and *ccr4*Δ::*gcn2*Δ mutants in either YPD or YNB-2% dextrose after 30 min at 37°C.

The reduction in *RPL2* at 37°C that was observed in the *ccr4*Δ mutant grown on minimal medium (**Fig 5B** center, **Fig 4C**) was abolished in the *ccr4*Δ::*gcn2*Δ mutant (**Fig 5B** right), suggesting that Gcn2 mediates the repression of RP mRNA in *C. neoformans* grown in minimal medium. Accordingly, there was no evidence of polysome collapse in the *ccr4*Δ::*gcn2*Δ double mutant grown in YNB minimal medium at 37°C (**Fig 5C**, right), which was observed as a larger 60S peak and concomitant smaller 80S peak in the *ccr4*Δ mutant in response to 37°C in the same medium (**Fig 5C**, center). This suggests that although Gcn2 is dispensable for temperature adaptation in YPD, Gcn2-dependent eIF2α phosphorylation is driving the translational response of the *ccr4*Δ mutant to temperature stress in minimal medium. This may indicate that growth in minimal medium is perceived as a translational stressor that when compounded with temperature leads to Gcn2 activation. These data also suggest that Gcn2 activation can compensate for the loss of Ccr4 in minimal medium with regard to translational repression (i.e., ribosome collapse and RP clearance), but this is not sufficient to maintain growth.

### Translational stress induces the ISR

In addition to the repression of cap-dependent translation initiation, Gcn2-mediated eIF2α phosphorylation in yeast promotes the translation of the transcription factor Gcn4, inducing the ISR [9]. The ISR regulon, defined as the transcriptional targets of Gcn4, have been characterized in *Saccharomyces cerevisiae*, and in filamentous ascomycetes as the cross-pathway control (*cpcA, cpc-1*) regulon [50].

To determine which stress triggers the ISR in *C. neoformans,* we measured protein levels of Gcn4, the expression of which is canonically upregulated when eIF2α is phosphorylated [11]. We expected to see in an increase in expression under conditions that induced eIF2α phosphorylation (**Fig 6A** and **Fig 1A**), and thus the greatest induction in response to 2 mM H_2_O_2_ (**Fig 6A** and **Fig 1A**), However, Western blotting showed only a moderate increase in Gcn4 protein in response to 2 mM H_2_O_2_ (**Fig 6B**, arrow indicates the band for Gcn4). Furthermore, 37°C stress and 1 mM H_2_O_2_ similarly induced Gcn4 even though we saw much greater phosphorylation of eIF2α in response to the oxidative stress (**Fig 6A** and **Fig 1A**). Also, 1 mM H_2_O_2_ and 40 mM 3-AT induced similar levels of eIF2α phosphorylation (**Fig 6A** and **Fig 1A**), but 40 mM 3-AT resulted in a much greater increase in Gcn4 protein levels (**Fig 6B**). These observations indicate that only modest levels of eIF2α phosphorylation are sufficient to trigger Gcn4 translation, and that the amount of Gcn4 does not scale with the magnitude of eIF2α phosphorylation.

**Fig 6.**
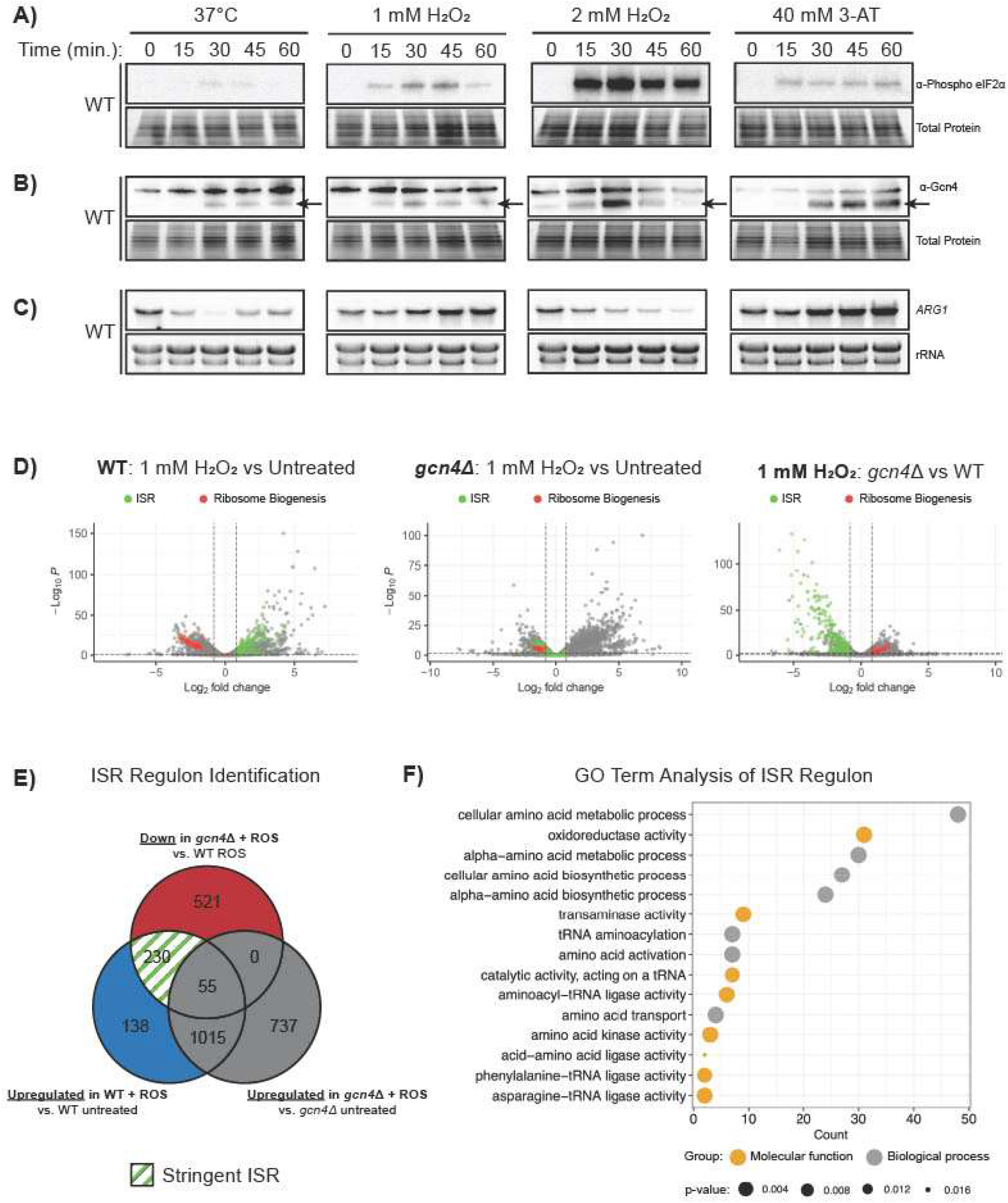
Temperature and oxidative stresses result in differential ISR induction. (**A**) Western blots for Gcn2 in the WT strain in response to 37°C, 1 mM H_2_O_2_, 2 mM H_2_O_2_, and 40 mM 3-AT. (**B**) Western blots for Gcn4 under the same conditions as panel. Arrows indicate the band for Gcn4.A (**C**) Northern blot analysis for the *ARG1* transcript under the same conditions as for panel A and B. (**D**) Volcano plots of –Log_10_P values versus Log_2_ fold change for RNA levels in the WT strain (1 mM H_2_O_2_ treated vs untreated; left), *gcn4*Δ mutant (1 mM H_2_O_2_ treated vs untreated; center), and the *gcn4*Δ mutant and WT (1 mM H_2_O_2_; right). Genes identified as part of the ISR are in green and ribosome biogenesis genes are in red. (**D**) Venn diagram describing the rationale used to determine the stringent ISR regulon. (**F**) GO enrichment analysis of genes identified as components of the ISR regulon.

To further assess ISR induction, we examined the expression pattern of *ARG1*, which is a known target of Gcn4 in *Saccharomyces cerevisiae* [12]. After confirming that *ARG1* is a bona fide target of Gcn4 in *C. neoformans* using the *gcn4Δ* mutant (**Fig S1**), we measured *ARG1* expression during 60 min of exposure to each stress. Northern blotting revealed that 1 mM H_2_O_2_ and 40 mM 3-AT induced *ARG1* expression, 2 mM H_2_O_2_ suppressed expression, but 37°C suppressed expression only transiently (**Fig 6C**).

Because the ISR is currently undescribed in *C. neoformans,* we performed RNA sequencing to identify the transcriptome in response to 60 min of oxidative stress (1 mM H_2_O_2_) in wild type and *gcn4Δ.* We initially defined the ISR regulon as transcripts that were downregulated (fold change [FC] < 1.75, *p* < 0.05) in the *gcn4Δ* mutant compared to WT in the context of oxidative stress (806 transcripts) (**Fig 6E**). However, this returned a substantial number of transcripts that either were not upregulated by the oxidative stress response or were upregulated even in the absence of Gcn4. We employed a more stringent analysis by focusing only on the transcripts that Gcn4-dependent up-regulation in the WT strain in response to oxidative stress (230 transcripts), which we defined as the *C. neoformans* ISR regulon (**Fig 6E**). A gene ontology (GO) term analysis was then performed on the 230 transcripts comprising the ISR regulon, revealing that the ISR primarily comprises components of amino acid biosynthetic pathways and tRNA regulation (**Fig 6F).**

Taken together, our observations demonstrate that the ISR in *C. neoformans* is induced by various host-relevant stresses as seen by Gcn4 translation, but the induction of one of its downstream targets, *ARG1*, varies according to the degree and type of stress. We also defined the *C. neoformans* ISR regulon, which closely matches that seen in *S. cerevisiae* literature with respect to the regulation of amino acid and tRNA metabolism [12].

## DISCUSSION

Part of the success of *C. neoformans* as a pathogen is its ability to respond and adapt to the environment inside the human lung. Although it is known that this requires mRNA decay-dependent translatome reprogramming [3,4], the mechanisms regulating this are unclear. We provide evidence supporting the roles of Ccr4-dependent mRNA decay and Gcn2-dependent eIF2α phosphorylation.

Our data demonstrate that temperature stress and various degrees of oxidative stress are sensed differently by *C. neoformans* and have different outcomes. Temperature stress resulted in rapid clearance of RP mRNAs and modest eIF2α phosphorylation, corresponding to a distinctly observable increase in collided ribosomes. Mild oxidative stress (from 1 mM H_2_O_2_) resulted in minimal RP mRNA repression and notable eIF2α phosphorylation without an increase in ribosome collisions, despite previous literature tying Gcn2 activation to ribosome collisions [21,25,51]. Interestingly, temperature stress and mild oxidative stress produced similar polysome profiles. These data, when viewed in the context of recent literature [17,52], suggest that temperature stress and mild oxidative stress result in differing states of stalled or collided ribosomes. Recent studies reveal that differing occupancy of tRNA binding sites on the ribosome results in the activation of different downstream quality control pathways [17,52], and specifically, an empty A site is required for Gcn2 activation [17]. This would suggest that the collided ribosomes as a result of temperature stress are stalled with an occupied A site rather than the empty A site required for Gcn2 activation.

We observed an association between ribosome collisions and accelerated RP mRNA repression. Recently, a role for Not5 in the link between ribosome collisions and mRNA surveillance and decay pathways was described, where Not5 interacts with a transiently unoccupied ribosome E site and recruits the Ccr4-NOT complex [52]. Our data demonstrate that the translational response to temperature is driven mainly by Ccr4-dependent mRNA decay, whereas the response to oxidative stress relies more heavily on Gcn2 activation. Thus, temperature stress may lead to ribosome collisions resulting in ribosomes with unoccupied E sites and accelerated Ccr4-mediated mRNA decay, whereas mild oxidative stress results in stalled ribosomes with unoccupied A sites, leading to increased Gcn2 activation.

Severe oxidative stress (from 2 mM H_2_O_2_) promoted RP mRNA decay and translational repression by a mechanism different from that induced by temperature and mild oxidative stress, possibly one that is uncoordinated by the cell and is instead a result of cellular damage. Oxidation of mRNAs can lead to decoding errors, and oxidation of ribosomal proteins and rRNA may result in dysfunctional ribosomes. Our observed lack of induction of *ARG1* in response to 2 mM H_2_O_2_ may even suggest that sufficient DNA damage precluded the transcriptional activation and translation of Gcn4. Nevertheless, *C. neoformans* can survive this stress, with no loss of viability on spot plates. This suggests that as a highly adaptive human pathogen, *C. neoformans* has multiple pathways to sense, respond to, and detoxify reactive oxygen species.

Growth in minimal medium exacerbates oxidative stress phenotypes while mitigating the responses to temperature stress. The mechanism responsible for this involves Gcn2, because the loss of *gcn2* in the double mutant reversed the loss of translational repression observed in the *ccr4*Δ mutant at 37°C in minimal medium. Thus, we propose an eIF2α phosphorylation “stress threshold” hypothesis, where cells grown in minimal medium are closer to the threshold of Gcn2 activation. These cells are then easily pushed over the threshold by temperature stress, which triggers the protection afforded by translational repression and ISR induction. Because of the stress from obligate anabolism in minimal medium, the cells express mRNAs encoding enzymes required for anabolism, which can be longer than RP mRNAs and more readily trigger translational pausing.

The recent identification of Gcn4 in *C. neoformans* enabled us to begin studying the role and regulation of the ISR [11]. Gcn2 activity was required for the translation of Gcn4, providing the first evidence, to our knowledge, of this conserved translational regulatory mechanism in basidiomycete fungi. Gcn4 translation did not necessarily correspond to eIF2α phosphorylation, though the modest levels of phospho-eIF2α induced by temperature and mild oxidative (1 mM H_2_O_2_) stresses were sufficient. This is consistent with the model in which phosphorylated eIF2α limits the availability of the 43S pre-initiation complex and promotes the bypass of the upstream open reading frame of the mRNA for Gcn4 [11]. Under severe oxidative stress (from 2 mM H_2_O_2_), when the levels of Gcn4 and eIF2α phosphorylation were highest, the transcription of ISR effector *ARG1* was repressed rather than induced. This suggests that there is a maximum level by which cells can cope with stress through the ISR, likely as a result of reduced transcription following this level of oxidative insult. Finally, the ISR regulon comprises a part of the translatome reprogramming in which translational resources are directed to stress-responsive mRNAs.

Together, these data support a model in which *C. neoformans* senses stress by monitoring its translational state and reprograms its translatome to combat the encountered stress. The differential responses of *C. neoformans* to temperature and oxidative stresses are particularly important because this organism is an environmental fungus, and the ability to survive at higher temperatures is crucial to the pathogenicity of this and other fungi to humans and other animals. Moreover, increasing global surface temperatures are likely to select for thermotolerance in other environmental pathogens. It is imperative that future work determines the molecular mechanisms that sense temperature stress and defines those responsible for promoting the stress-responsive translatome.

## DATA AVAILABILITY

The sequencing data can be viewed at GEO under accession number: GSE206508 with the reviewer token: uritkmwqbxclbin

## ACKNOWLEDGMENTS

We would like to thank Dr. Karen Dietz for assistance in editing this manuscript. The *gcn4*Δ strain was obtained from the Fungal Genetics Stock Center (Manhattan, Kansas, USA) as part of the Madhani *Cryptococcus neoformans* deletion collection.

## REFRENCES

1. Rajasingham, R.; Smith, R.M.; Park, B.J.; Jarvis, J.N.; Govender, N.; Chiller, T.M.; Denning, D.W.; Loyse, A.; Boulware, D.R. Global burden of disease of HIV-associated cryptococcal meningitis: an updated analysis. Lancet Infect. Dis. 2017, 17, 873–881, doi:10.1016/j.celrep.2015.11.047.Long.

2. Bloom, A.L.M.; Goich, D.; Knowles, C.M.; Panepinto, J.C. Glucan Unmasking Identi fi es Regulators of Temperature-Induced Translatome Reprogramming in C. neoformans. 2021, 6, doi:10.1128/mSphere.01281-20.

3. Bloom, A.L.M.; Jin, R.M.; Leipheimer, J.; Bard, J.E.; Yergeau, D.; Wohlfert, E.A.; Panepinto, J.C. Thermotolerance in the pathogen Cryptococcus neoformans is linked to antigen masking via mRNA decay-dependent reprogramming. Nat. Commun. 2019, 10, 4950, doi:10.1038/s41467-019-12907-x.

4. Bloom, A.L.M.; Solomons, J.T.G.; Havel, V.E.; Panepinto, J.C. Uncoupling of mRNA synthesis and degradation impairs adaptation to host temperature in Cryptococcus neoformans. Mol. Microbiol. 2013, 89, 65–83, doi:10.1111/mmi.12258.

5. Leipheimer, J.; Bloom, A.L.M.; Campomizzi, C.S.; Salei, Y.; Panepinto, J.C. Translational Regulation Promotes Oxidative Stress Resistance in the Human Fungal Pathogen Cryptococcus neoformans. MBio 2019, 10, 8–11, doi:10.1128/mBio.02143-19.

6. Ho, Y.H.; Shishkova, E.; Hose, J.; Coon, J.J.; Gasch, A.P. Decoupling Yeast Cell Division and Stress Defense Implicates mRNA Repression in Translational Reallocation during Stress. Curr. Biol. 2018, 28, 2673–2680.e4, doi:10.1016/j.cub.2018.06.044.

7. Sonenberg, N.; Hinnebusch, A.G. Regulation of Translation Initiation in Eukaryotes: Mechanisms and Biological Targets. Cell 2009, 136, 731–745, doi:10.1016/j.cell.2009.01.042.

8. Cao, D.; Parker, R. Computational modeling of eukaryotic mRNA turnover. Rna 2001, 7, 1192–1212, doi:10.1017/S1355838201010330.

9. Dever, T.E.; Feng, L.; Wek, R.C.; Cigan, A.M.; Donahue, T.F.; Hinnebusch, A.G. Phosphorylation of initiation factor 2α by protein kinase GCN2 mediates genespecific translational control of GCN4 in yeast. Cell 1992, 68, 585–596, doi:10.1016/0092-8674(92)90193-G.

10. Hinnebusch, A.G. TRANSLATIONAL REGULATION OF GCN4 AND THE GENERAL AMINO ACID CONTROL OF YEAST. Annu. Rev. Microbiol. 2005, 59, 407–450, doi:10.1146/annurev.micro.59.031805.133833.

11. Wallace, E.W.J.; Maufrais, C.; Sales-Lee, J.; Tuck, L.R.; de Oliveira, L.; Feuerbach, F.; Moyrand, F.; Natarajan, P.; Madhani, H.D.; Janbon, G. Quantitative global studies reveal differential translational control by start codon context across the fungal kingdom. Nucleic Acids Res. 2020, 48, 2312–2331, doi:10.1093/nar/gkaa060.

12. Rawal, Y.; Chereji, R. V.; Valabhoju, V.; Qiu, H.; Ocampo, J.; Clark, D.J.; Hinnebusch, A.G. Gcn4 Binding in Coding Regions Can Activate Internal and Canonical 5’ Promoters in Yeast. Mol. Cell 2018, 70, 297–311.e4, doi:10.1016/j.molcel.2018.03.007.

13. Jobava, R.; Mao, Y.; Guan, B.J.; Hu, D.; Krokowski, D.; Chen, C.W.; Shu, X.E.; Chukwurah, E.; Wu, J.; Gao, Z.; et al. Adaptive translational pausing is a hallmark of the cellular response to severe environmental stress. Mol. Cell 2021, 81, 4191–4208.e8, doi:10.1016/j.molcel.2021.09.029.

14. Liaud, N.; Horlbeck, M.A.; Gilbert, L.A.; Gjoni, K.; Weissman, J.S.; Cate, J.H.D. Cellular response to small molecules that selectively stall protein synthesis by the ribosome. PLoS Genet. 2019, 15, 1–30, doi:10.1371/journal.pgen.1008057.

15. Simms, C.L.; Hudson, B.H.; Mosior, J.W.; Rangwala, A.S.; Zaher, H.S. An Active Role for the Ribosome in Determining the Fate of Oxidized mRNA. Cell Rep. 2014, 9, 1256–1264, doi:10.1016/j.celrep.2014.10.042.

16. Rubio, A.; Ghosh, S.; Mülleder, M.; Ralser, M.; Mata, J. Ribosome profiling reveals ribosome stalling on tryptophan codons and ribosome queuing upon oxidative stress in fission yeast. Nucleic Acids Res. 2021, 49, 383–399, doi:10.1093/nar/gkaa1180.

17. Yan, L.L.; Zaher, H.S. Ribosome quality control antagonizes the activation of the integrated stress response on colliding ribosomes. Mol. Cell 2021, 81, 614–628.e4, doi:10.1016/j.molcel.2020.11.033.

18. Wu, C.C.C.; Zinshteyn, B.; Wehner, K.A.; Green, R. High-Resolution Ribosome Profiling Defines Discrete Ribosome Elongation States and Translational Regulation during Cellular Stress. Mol. Cell 2019, 73, 959–970.e5, doi:10.1016/j.molcel.2018.12.009.

19. Merret, R.; Nagarajan, V.K.; Carpentier, M.C.; Park, S.; Favory, J.J.; Descombin, J.; Picart, C.; Charng, Y.Y.; Green, P.J.; Deragon, J.M.; et al. Heat-induced ribosome pausing triggers mRNA co-translational decay in Arabidopsis thaliana. Nucleic Acids Res. 2015, 43, 4121–4132, doi:10.1093/nar/gkv234.

20. Shalgi, R.; Hurt, J.A.; Krykbaeva, I.; Taipale, M.; Lindquist, S.; Burge, C.B. Widespread Regulation of Translation by Elongation Pausing in Heat Shock. Mol. Cell 2013, 49, 439–452, doi:10.1016/j.molcel.2012.11.028.

21. Wu, C.C.; Peterson, A.; Zinshteyn, B.; Regot, S.; Green, R. Ribosome Collisions Trigger General Stress Responses to Regulate Cell Fate. Cell 2020, 182, 404–416.e14, doi:10.1016/j.cell.2020.06.006.

22. Darnell, A.M.; Subramaniam, A.R.; O’Shea, E.K. Translational Control through Differential Ribosome Pausing during Amino Acid Limitation in Mammalian Cells. Mol. Cell 2018, 71, 229–243.e11, doi:10.1016/j.molcel.2018.06.041.

23. Yan, L.L.; Zaher, H.S. Ribosome quality control antagonizes the activation of the integrated stress response on colliding ribosomes. Mol. Cell 2020, 1–15, doi:10.1016/j.molcel.2020.11.033.

24. Inglis, A.J.; Masson, G.R.; Shao, S.; Perisic, O.; McLaughlin, S.H.; Hegde, R.S.; Williams, R.L. Activation of GCN2 by the ribosomal P-stalk. Proc. Natl. Acad. Sci. U. S. A. 2019, 116, 4946–4954, doi:10.1073/pnas.1813352116.

25. Ishimura, R.; Nagy, G.; Dotu, I.; Chuang, J.H.; Ackerman, S.L. Activation of GCN2 kinase by ribosome stalling links translation elongation with translation initiation. Elife 2016, 5, 1–22, doi:10.7554/eLife.14295.

26. Hickey, K.L.; Dickson, K.; Cogan, J.Z.; Replogle, J.M.; Schoof, M.; D’Orazio, K.N.; Sinha, N.K.; Hussmann, J.A.; Jost, M.; Frost, A.; et al. GIGYF2 and 4EHP Inhibit Translation Initiation of Defective Messenger RNAs to Assist Ribosome-Associated Quality Control. Mol. Cell 2020, 79, 950–962.e6, doi:10.1016/j.molcel.2020.07.007.

27. Guydosh, N.R.; Green, R. Dom34 rescues ribosomes in 3’ untranslated regions. Cell 2014, 156, 950–962, doi:10.1016/j.cell.2014.02.006.

28. Ikeuchi, K.; Inada, T. Ribosome-associated Asc1/RACK1 is required for endonucleolytic cleavage induced by stalled ribosome at the 3’ end of nonstop mRNA. Sci. Rep. 2016, 6, 1–10, doi:10.1038/srep28234.

29. Juszkiewicz, S.; Chandrasekaran, V.; Lin, Z.; Kraatz, S.; Ramakrishnan, V.; Hegde, R.S. ZNF598 Is a Quality Control Sensor of Collided Ribosomes. Mol. Cell 2018, 72, 469–481.e7, doi:10.1016/j.molcel.2018.08.037.

30. Doma, M.K.; Parker, R. Endonucleolytic cleavage of eukaryotic mRNAs with stalls in translation elongation. Nature 2006, 440, 561–564, doi:10.1038/nature04530.

31. Simms, C.L.; Yan, L.L.; Zaher, H.S. Ribosome Collision Is Critical for Quality Control during No-Go Decay. Mol. Cell 2017, 68, 361–373.e5, doi:10.1016/j.molcel.2017.08.019.

32. D’Orazio, K.N.; Wu, C.C.C.; Sinha, N.; Loll-Krippleber, R.; Brown, G.W.; Green, R. The endonuclease Cue2 cleaves mRNAs at stalled ribosomes during no go decay. Elife 2019, 8, 1–27, doi:10.7554/eLife.49117.001.

33. Matsuo, Y.; Ikeuchi, K.; Saeki, Y.; Iwasaki, S.; Schmidt, C.; Udagawa, T.; Sato, F.; Tsuchiya, H.; Becker, T.; Tanaka, K.; et al. Ubiquitination of stalled ribosome triggers ribosome-associated quality control. Nat. Commun. 2017, 8, 1–13, doi:10.1038/s41467-017-00188-1.

34. Ikeuchi, K.; Tesina, P.; Matsuo, Y.; Sugiyama, T.; Cheng, J.; Saeki, Y.; Tanaka, K.; Becker, T.; Beckmann, R.; Inada, T. Collided ribosomes form a unique structural interface to induce Hel2-driven quality control pathways. EMBO J. 2019, 38, 1–21, doi:10.15252/embj.2018100276.

35. Kostova, K.K.; Hickey, K.L.; Osuna, B.A.; Hussmann, J.A.; Frost, A.; Weinberg, D.E.; Weissman, J.S. CAT-tailing as a fail-safe mechanism for efficient degradation of stalled nascent polypeptides. Science (80-.). 2017, 357, 414–417, doi:10.1126/science.aam7787.

36. Brandman, O.; Stewart-Ornstein, J.; Wong, D.; Larson, A.; Williams, C.C.; Li, G.W.; Zhou, S.; King, D.; Shen, P.S.; Weibezahn, J.; et al. A Ribosome-Bound Quality Control Complex Triggers Degradation of Nascent Peptides and Signals Translation Stress. Cell 2012, 151, 1042–1054, doi:10.1016/j.cell.2012.10.044.

37. Panepinto, J.; Liu, L.; Ramos, J.; Zhu, X.; Valyi-Nagy, T.; Eksi, S.; Fu, J.; Jaffe, H.A.; Wickes, B.; Williamson, P.R. The DEAD-box RNA helicase Vad1 regulates multiple virulence-associated genes in Cryptococcus neoformans. J. Clin. Invest. 2005, 115, 632–641, doi:10.1172/JCI200523048.

38. Panepinto, J.C.; Oliver, B.G.; Amlung, T.W.; Askew, D.S.; Rhodes, J.C. Expression of the Aspergillus fumigatus rheb homologue, rhbA, is induced by nitrogen starvation. Fungal Genet. Biol. 2002, 36, 207–214, doi:10.1016/S1087-1845(02)00022-1.

39. Martin, M. Cutadapt removes adapter sequences from high-throughput sequencing reads. EMBnet.journal 2011, 17, 10, doi:10.14806/ej.17.1.200.

40. Dobin, A.; Davis, C.A.; Schlesinger, F.; Drenkow, J.; Zaleski, C.; Jha, S.; Batut, P.; Chaisson, M.; Gingeras, T.R. STAR: ultrafast universal RNA-seq aligner. Bioinformatics 2013, 29, 15–21, doi:10.1093/bioinformatics/bts635.

41. Parrish, N.; Hormozdiari, F.; Eskin, E. RSEM: accurate transcript quantification from RNA-Seq data with or without a reference genome. Bioinforma. Impact Accurate Quantif. Proteomic Genet. Anal. Res. 2014, 21–40, doi:10.1201/b16589.

42. Love, M.I.; Huber, W.; Anders, S. Moderated estimation of fold change and dispersion for RNA-seq data with DESeq2. Genome Biol. 2014, 15, 550, doi:10.1186/s13059-014-0550-8.

43. Blighe K; Rana S; Lewis M Enhanced Volcano: Publication-ready volcano plots with enhanced colouring and labeling 2022.

44. Wickham, H. ggplot2; Springer New York: New York, NY, 2009; ISBN 978-0-387-98140-6.

45. Wu, C.C.-C.; Zinshteyn, B.; Wehner, K.A.; Green, R. High-Resolution Ribosome Profiling Defines Discrete Ribosome Elongation States and Translational Regulation during Cellular Stress. Mol. Cell 2019, 73, 959–970.e5, doi:10.1016/j.molcel.2018.12.009.

46. Bloom, A.L.M.; Leipheimer, J.; Panepinto, J.C. mRNA decay: an adaptation tool for the environmental fungal pathogen Cryptococcus neoformans. Wiley Interdiscip. Rev. RNA 2017, 8, e1424, doi:10.1002/wrna.1424.

47. Glazier, V.E.; Panepinto, J.C. The ER stress response and host temperature adaptation in the human fungal pathogen Cryptococcus neoformans. Virulence 2014, 5, 351–356.

48. Banerjee, D.; Bloom, A.L.M.; Panepinto, J.C. Opposing PKA and Hog1 signals control the post-transcriptional response to glucose availability in Cryptococcus neoformans. Mol. Microbiol. 2016, 102, 306–320, doi:10.1111/mmi.13461.

49. Panepinto, J.C.; Komperda, K.W.; Hacham, M.; Shin, S.; Liu, X.; Williamson, P.R. Binding of Serum Mannan Binding Lectin to a Cell Integrity-Defective Cryptococcus neoformans ccr4Δ Mutant. Infect. Immun. 2007, 75, 4769–4779, doi:10.1128/IAI.00536-07.

50. Carsiotis, M.; Jones, R.F. Cross-Pathway Regulation: Tryptophan-Mediated Control of Histidine and Arginine Biosynthetic Enzymes in Neurospora crassa. J. Bacteriol. 1974, 119, 889–892, doi:10.1128/jb.119.3.889-892.1974.

51. Pochopien, A.A.; Beckert, B.; Kasvandik, S.; Berninghausen, O.; Beckmann, R.; Tenson, T.; Wilson, D.N. Structure of Gcn1 bound to stalled and colliding 80S ribosomes. Proc. Natl. Acad. Sci. U. S. A. 2021, 118, doi:10.1073/pnas.2022756118.

52. Buschauer, R.; Matsuo, Y.; Sugiyama, T.; Chen, Y.H.; Alhusaini, N.; Sweet, T.; Ikeuchi, K.; Cheng, J.; Matsuki, Y.; Nobuta, R.; et al. The Ccr4-Not complex monitors the translating ribosome for codon optimality. Science (80-.). 2020, 368, doi:10.1126/science.aay6912.

