## Supplemental Figures for "Contributions of Ccr4 and Gcn2 to the translational response of *C. neoformans* to host-relevant stressors and Integrated Stress Response induction"

### Figure S1:

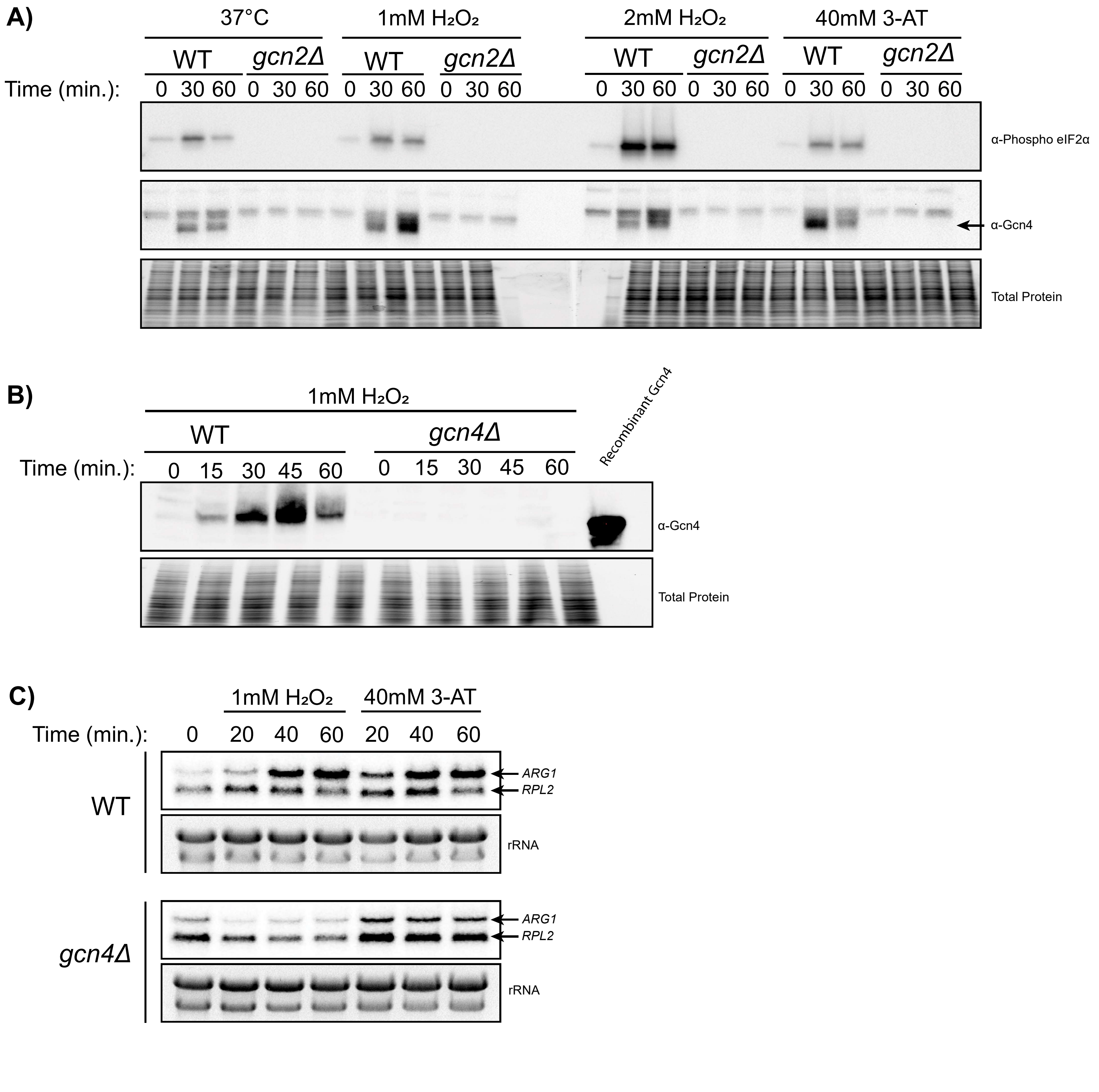
A)
37°C
1mM H₂O₂
2mM H₂O₂
40mM 3-AT
WT
gcn2Δ
Time (min.):
0 30 60
α-Phospho eIF2α
α-Gcn4
Total Protein
B)
1mM H₂O₂
WT
gcn4Δ
Time (min.):
0 15 30 45 60
Recombinant Gcn4
α-Gcn4
Total Protein
C)
1mM H₂O₂
40mM 3-AT
Time (min.):
0 20 40 60 20 40 60
WT
gcn4Δ
ARG1
RPL2
rRNA

**SUPPLEMENTAL MATERIAL**

**Table S1. Primers**

| **Primer name** | **Sequence** |
|---|---|
| F-NEO-*Bgl*II | TAATAAAGATCTAGCGGATAACAATTTCACACAGG |
| R-NEO-*Sac*I | TAATAACAGCTCCGACGGCCAGTGAATTGTAATACG |
| F-CCR4-1kbUp-Spe1 | TAATAAACTAGTCTGCTGTTTCAACTCCATAGGC |
| R-CCR4-1kbDown-Spe1 | TAATAAACTAGTGGAATAGTTTGACGGGTGG |
| ARG1-Northern-F | GGTGCTGTCTTCCATCTTG |
| ARG1-Northern-R | ACAGTCTTTTGAGATGCGGGGA |

**Fig S1.** (**A**) Western blots for eIF2α phosphorylation (top) and Gcn4 (middle) in the WT and *gcn2*Δ strains in response to 37°C, 1 mM $H_2O_2$, 2 mM $H_2O_2$, and 40 mM 3-AT. Arrow indicates band for Gcn4. (**B**) Western blots for Gcn4 in response to 1 mM $H_2O_2$ in the WT and *gcn4*Δ strains. Recombinant Gcn4 protein was included for antibody validation. (**C**) Northern blot analysis for the *ARG1* and *RPL2* transcripts in response to 1 mM $H_2O_2$ and 40 mM 3-AT in the WT and *gcn4*Δ strains.
